# Lauric acid: a promising antimicrobial for the selective inhibition of *Staphylococcus epidermidis* strains associated with infection

**DOI:** 10.1101/2025.10.06.680683

**Authors:** Elisabete Morais, Filipe Magalhães, Ons Bouchami, Dragana PC de Barros, Luís G Gonçalves, Abel Oliva, Maria Miragaia, Ana V Coelho

## Abstract

*Staphylococcus epidermidis*, a human skin colonizer and opportunistic pathogen associated with medical device infections, comprises two phylogenetic lineages: A/C (infection and colonization strains) and B (primarily colonization strains). Given the need for antibiotic alternatives, we investigated the antimicrobial activity of skin-derived fatty acids—lauric, palmitoleic, and linoleic acids—against representative strains from both lineages. Fatty acids reduced exponential growth rate and maximum population, with greater effects on the A/C strain. Lauric and palmitoleic acids decreased colony radius in the B strain. Importantly, lauric acid showed no cytotoxicity on a 3D reconstructed human epidermis model. Our findings demonstrate differential susceptibility between lineages, with A/C strains showing greater sensitivity to all tested fatty acids than B strains. These results highlight the potential of lauric acid as a topical formulation for selective inhibition of *S. epidermidis* growth, offering a promising approach for preventing pathogenic strain proliferation while preserving beneficial skin colonizers.

## 1. Introduction

Healthcare-associated infections pose a major global health challenge, affecting 3.2% and 6.5% of hospitalized patients in the United States and European Union, respectively, largely due to rising antibiotic resistance [1,2].

*Staphylococcus epidermidis* (SE), the most abundant skin symbiont, has emerged as an opportunistic nosocomial pathogen [3] and the leading cause of medical device-associated infections [4], particularly in neonates with increasing prevalence [5,6]. In the more drastic scenarios, SE infection leads to severe morbidity or death [5]. SE comprises two main clonal lineages with distinct pathogenic potential: A/C lineage exhibits broader skin niche occupation, increased antibiotic resistance, and enhanced biofilm production at blood pH, while B lineage is adapted to microaerobic, lipid-rich environments of hair follicles and sebaceous glands [7,8].

Fatty acids (FAs) represent key antimicrobial components of skin’s innate immunity, synthesized in the epidermis or secreted by sebaceous glands. They act as antimicrobial agents and confer protection against pathogens [12]. As promising antibiotic alternatives, they demonstrate potent activity against multidrug-resistant strains [9–11]. Moreover, they facilitate antibiotic penetration when co-administered [10] while causing no skin irritation upon topical application [9]

Among tested FAs, lauric acid (LA, C12:0) showed strong antimicrobial effects against both SE and S. *aureus* [11,12], while palmitoleic acid (PA, C16:1) and linoleic acid (LIN, C18:2) exhibit reduced activity against SE [11–14]. This suggests that staphylococci use diverse mechanisms to respond to endogenous FAs. Importantly, strain-level sensitivity variations suggest that FAs could serve as selective antimicrobials. Since commensal SE strains benefit host skin health [15], antimicrobials should preferentially target pathogenic strains. Lineage B strains possess operons for staphyloxanthin biosynthesis and ESAT-6 secretion that confer FA resistance [7,16,17], suggesting that endogenous FAs could selectively target pathogenic A/C strains for treating/preventing device-related infections [18,19].

This study evaluated the antimicrobial effectiveness of three FAs against representative SE strains: ICE25 (lineage A/C, infection isolate) and 19N (lineage B, healthy donor isolate).

## 2. Materials and Methods

### 2.1. Study Design

The antimicrobial effects of three FA (LA, LIN, and PA) were evaluated against *Staphylococcus epidermidis* (SE) using multiple growth conditions: liquid media, solid agar, and a 3D reconstructed human epidermis (RHE) model. Minimum inhibitory concentrations (MIC) were determined, growth curves analysed, and cytotoxicity assessed to identify non-toxic concentrations for skin application studies. Detailed protocols are provided as Supplementary Materials.

### 2.2. Bacterial Strains and Culture Conditions

Representative SE strains from phylogenetic lineages B (19N) and A/C (ICE25) (https://www.ncbi.nlm.nih.gov/bioproject/?term=PRJNA606771) were used. Cultures were grown on tryptic soy agar (TSA) and in tryptic soy broth (TSB) at 37°C and pH 5.5, adjusted with HCl before sterilization. All bacterial assays were performed under these standardized conditions.

### 2.3. Fatty Acid Preparation

Stock solutions of LA, PA, and LIN (Table S1) were prepared fresh daily in absolute ethanol and diluted in appropriate media supplemented with 0.1% DMSO. For cytotoxicity assays and assays with RHE, FA stock solutions were diluted in phosphate-buffered saline (PBS). Final ethanol concentrations were maintained at 0.5% (v/v) for bacterial assays and 5% for cytotoxicity studies.

### 2.4. Antimicrobial Activity Assessment

FA MIC values were determined using the broth microdilution method in TSB with an initial inoculum of 5×10^5^ CFU/well. After 20 hours at 37°C, growth inhibition was assessed by optical density measurements at 595 nm. Growth curves were generated using increasing FA concentrations in aerated TSB, to determine lag time and exponential growth rate. For solid media assays, 0.5×10^3^ CFU were plated on FA-supplemented TSA, and colony size and viability were assessed after 24-40 hours using optical microscopy and colony counting.

### 2.5. Cytotoxicity Evaluation

HaCaT keratinocytes were cultured in DMEM with 10% FBS and exposed to FA concentrations (MIC/2, MIC, 2×MIC) for 3 hours. Cell viability was determined using the MTT assay, measuring absorbance at 540 nm. For 3D studies, RHE was generated using neonatal human epidermal keratinocytes cultured at the air-liquid interface for 12 days until fully differentiated, following established protocol. Cytotoxicity on RHE was assessed according to OECD Test Guideline 439, with FA solutions applied topically for 3 hours, followed by MTT viability assessment.

### 2.6. Antibacterial Studies onto RHE

RHE was generated as described above. For antibacterial studies, RHE was colonized with 1×10^2^ CFU of each SE strain for 3 hours, then treated with RHE non-toxic LA concentrations (7.81 μg/mL) for 1h. Bacterial recovery was quantified by plating wash suspensions and calculating growth inhibition relative to controls.

### 2.7. Histological Analysis

RHE tissues were fixed in 10% neutral-buffered formalin, sectioned (5 mm), and stained with hematoxylin-eosin for morphological analysis using fluorescence microscopy and ImageJ software.

### 2.8. Statistical Analysis

Comparisons between strains and treatments were performed using *t*-tests for MIC and growth data, and Mood’s median test for colony size analysis. Statistical significance was set at p-value < 0.05. All analyses were conducted in Python 3.11.5 using custom scripts available at https://github.com/eccmorais/MIC_Fatty_acids_Sepidermidis.git.

## 3. Results

Given SE’s dual role as commensal and opportunistic pathogen, effective antimicrobials should ideally target infection-associated A/C lineage strains while preserving beneficial B lineage commensals. Here, we evaluated the antimicrobial activity of selected FAs against representative strains from both SE phylogenetic lineages.

### 3.1. A/C strain is more susceptible to LA, PA and LIN

Broth microdilution assays revealed that ICE25 (A/C lineage) was more susceptible to all tested FAs than 19N (B lineage) (Figure 1). LA demonstrated the strongest antimicrobial activity with MIC values of 7.81 µg/mL for ICE25 and 15.63 µg/mL for 19N. PA showed intermediate activity with an MIC of 31.25 µg/mL for ICE25, while 19N growth was inhibited by only 50% at this concentration. LIN exhibited weak antimicrobial effects, achieving maximum growth inhibition of 40% (ICE25) and 25% (19N) at 4000 µg/mL—over 200-fold higher than LA’s MIC.

**Figure 1.**
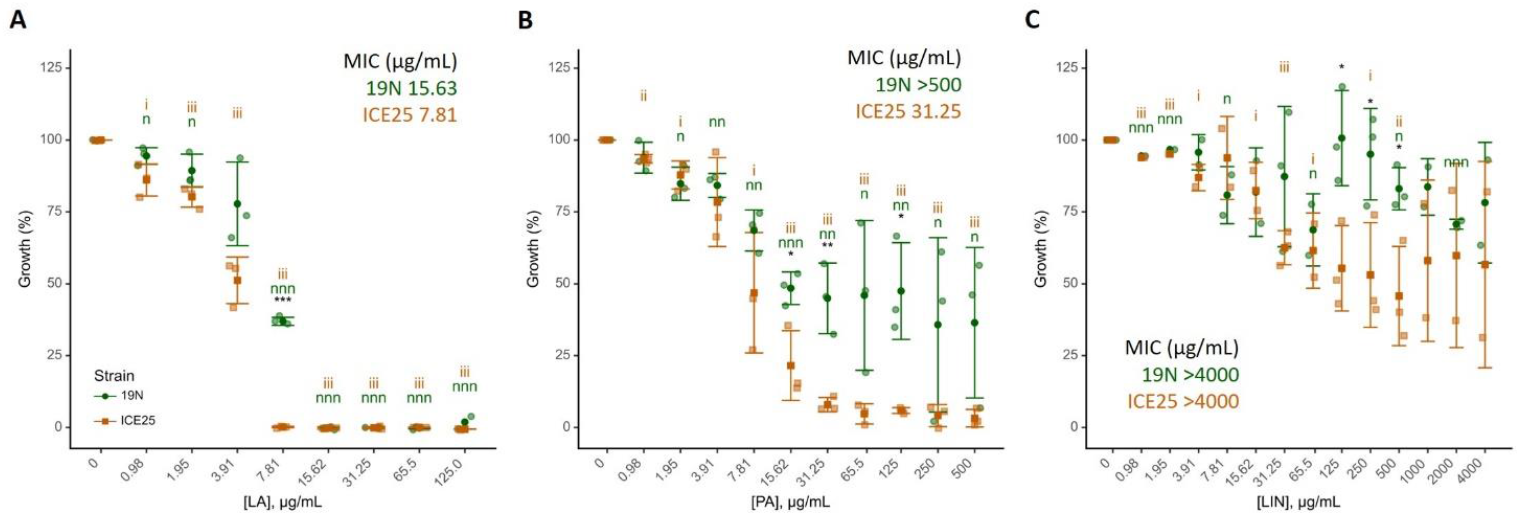
FA effect against SE. Percentage of growth, calculated from OD_595nm_ measurements in MIC determination by the broth microdilution method and considering concentration 0 µg/mL as 100% of growth, for 19N and ICE25 strains in the presence of LA (**A**), PA (**B**), and LIN (**C**) in TSB at pH 5.5. Values are represented for each biological replicate, average (darker symbol) and standard deviation (n=3). The x-axis is log2 transformed for visualisation purposes. *t*-test results are represented for comparison: 1) between strains for a certain experimental condition (*); and 2) within the 19N/ICE25 strains against the respective control (0 µg/mL FAs), n and i, for 19N and ICE25, respectively. ^***/nnn/iii^ *p*-value < 0.001; ^**/nn/ii^ *p*-value < 0.01, ^*/n/ i^ *p*-value < 0.05.

Strain-discriminative effects were observed at 7.81 µg/mL for LA, 15.63-125 µg/mL for PA, and 125-500 µg/mL for LIN. Variability in FA responses, particularly for 19N with PA and both strains with LIN, suggests cellular heterogeneity and diverse adaptation mechanisms within each strain. While LA had the lowest MIC against ICE25, PA demonstrated superior selective potential, completely inhibiting ICE25 growth across a broader concentration range (31.25-500 µg/mL) with minimal effect on 19N.

### 3.2. LA and PA mainly induce 19N lag time increase and ICE25 maximum population decrease

Growth assays using sub-MIC, MIC, and supra-MIC concentrations were performed in larger volumes (500-fold) under agitation to enhance oxygenation (Figures 2, 3; Table S2). These altered conditions may explain some discrepancies with the previous assays. High variation in lag phase duration was observed for LA and PA concentrations within each strain, while growth curves overlapped in FA absence, suggesting, as in the previous assay, that bacterial population heterogeneity plays an important role in SE adaptation to FAs.

**Figure 2.**
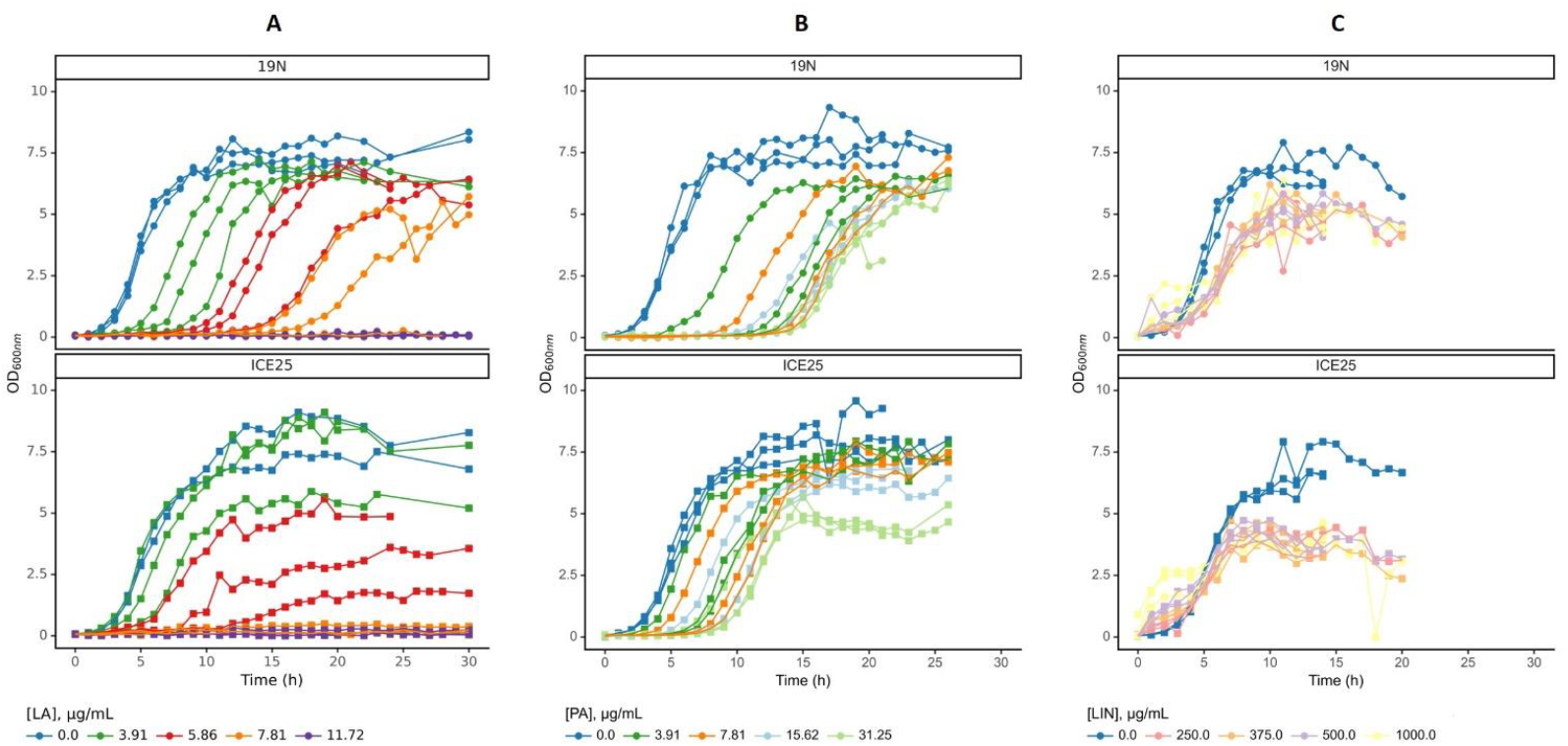
Growth curves of SE strains in the presence of LA, PA and LIN. 19N and ICE25 were grown in the presence of increasing concentrations of LA (**A**), PA (**B**), and LIN (**C**) in TSB pH 5.5. Individual growth curves are represented for each biological replicate (n=3), being those for the same FA concentration represented with the same colour.

**Figure 3.**
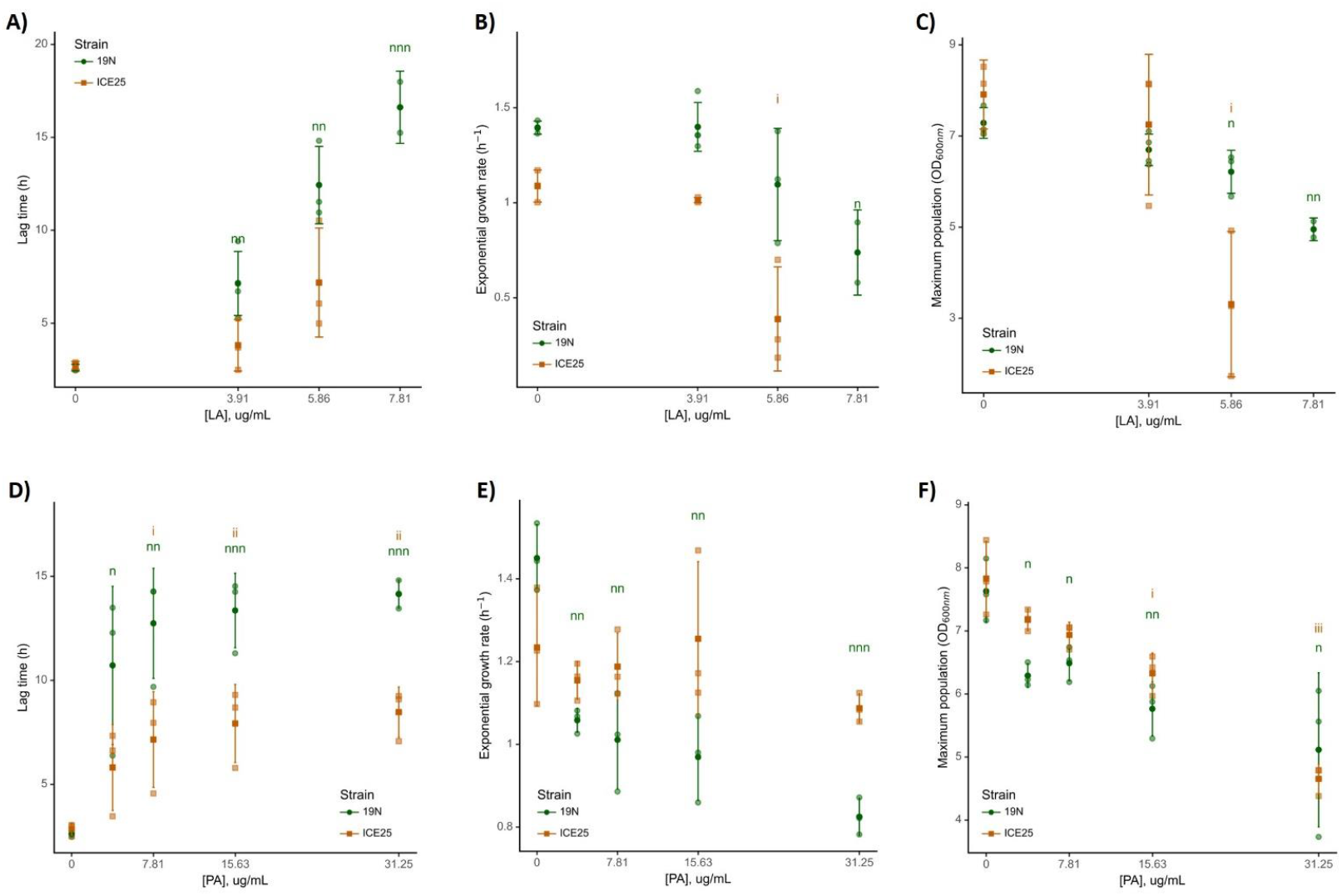
LA and PA effect on the SE growth parameters. 19N and ICE25 strains were grown in TSB at pH 5.5 with increasing concentrations of LA (**A, B, C**) and PA (**D, E, F**). Lag time (h), exponential growth rate (h^-1^), and maximum population (OD_600nm_) were calculated using the Curveball Python module (Ram 2019). Values are represented by biological replicas along average (dark point) and standard deviation (n=3). *t*-test results are represented for comparison within 19N/ICE25 strains against the respective control (0 µg/mL FAs), n and i, for 19N and ICE25, respectively. ^nnn/iii^ *p*-value < 0.001; ^nn/ii^ *p*-value < 0.01, ^n/ i^ *p*-value < 0.05.

LA completely inhibited both strains at 11.72 µg/mL, while at 7.81 µg/mL ICE25 showed no growth and 19N exhibited a 14-hour growth delay (Figure 2; Table S2a). LA significantly increased lag phase duration only in 19N, which is significantly different between the two strains at 5.86 µg/mL (Figure 3A). A significant decrease in the absolute exponential growth rate was determined for 7.81 µg/mL for 19N and 5.86 µg/mL for ICE25 (Figure 3B, Table S2a). A decrease in the absolute maximum population was observed for 19N at 5.86 µg/mL (Figure 3C, Table S2a), with ICE25 suffering a significantly higher decrease (Table S2b).

PA increased lag phase from 3.91 µg/mL (19N) and 7.81 µg/mL (ICE25), with 19N showing significantly longer delays from 7.81 µg/mL (Figure 3D, Table S2a). PA reduced absolute exponential growth rate and absolute maximum population in 19N from 3.91 µg/mL PA (Figure 3E/F, Table S2a). The exponential growth rate starts to be significantly different between the two strains at 7.81 µg/mL of PA (Figure 3E, Table S2b). ICE25 maximum population decreased only at 15.62 µg/mL (Figure 3F; Table S2a).

LIN produced optical density artefacts due to poor media dissolution, creating broad peaks during the first 4 hours that increased with concentration and prevented accurate growth parameter determination (Figure 2C, Figure S1). Despite this limitation, both strains showed reduced growth rates and population sizes, with greater effects on ICE25 (Figure 2C). Due to these technical issues, LIN was excluded from subsequent assays.

These results demonstrate LA’s selective antimicrobial potential against infection-associated SE strains at 6-8 µg/mL, while PA reduced maximum populations (40% for ICE25, 33% for 19N) without complete growth inhibition, contrasting with broth microdilution findings.

### 3.3. LA reduces more the number of A/C strain CFUs and its radius

To mimic skin conditions, CFU reduction was assessed on TSA at pH 5.5 (Figure 4; Table S3). The impact of increasing concentrations of LA and PA on the number of CFU relative to control at 40h was evaluated (Figure 4 A/B and Table S3). LA reduced colony numbers starting at 6.0 µg/mL and 7.81 µg/mL, respectively for 19N and ICE25. Complete ICE25 growth inhibition occurred at ≥10 µg/mL LA, while 19N retained ∼38% growth at 12 µg/mL (Figure 4A). PA treatment resulted in similar responses for both strains, with ∼15% growth remaining at 31.25 µg/mL (Figure 4B).

**Figure 4.**
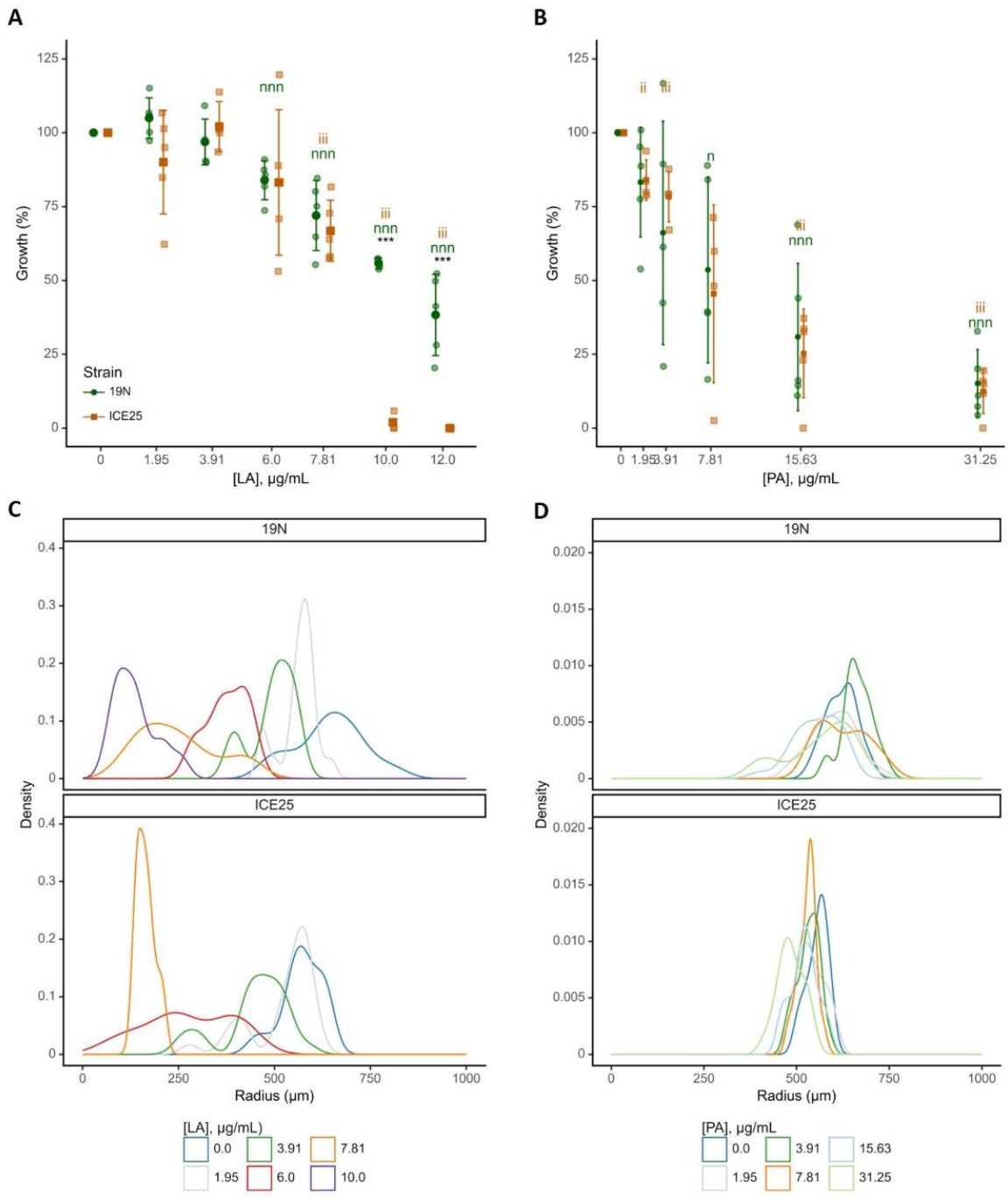
Effect of LA and PA on the number of CFUs and colony radius. Percentage of CFU after 40 hours of incubation, calculated from CFUs counted for several FA concentrations and considering concentration 0 µg/mL as 100% of growth, for 19N and ICE25 strains in the presence of LA (**A**) and PA (**B**) (n=5). *t*-test results are represented for comparison: 1) between strains for a certain experimental condition (*); and 2) within the 19N/ICE25 strains against the respective control (0 µg/mL FAs), n and i, for 19N and ICE25, respectively. ^***/nnn/iii^ *p*-value < 0.001; ^**/nn/ii^ *p*-value < 0.01, ^*/n/ i^ *p*-value < 0.05. Colonies size distribution after 24 hours of growth in TSA plates at pH 5.5 supplemented with LA (**C**) and PA (**D**) (n=25).

Colony radius measurements after 24 hours revealed heterogeneous size distributions with two predominant radii, with 19N showing larger median colony sizes in control conditions (Figures 4C/D; Table S3). LA induced concentration-dependent radius reduction while maintaining bimodal size distributions, with larger colonies predominating at lower concentrations and smaller colonies at ∼8 µg/mL (Figure 4C). PA showed less pronounced effects on colony size, though bimodal distributions persisted with greater radius differences in 19N. The maximum radius reduction occurred at 7.81 µg/mL PA for ICE25 (Figures 4D and Table S3).

These results confirm LA’s superior antimicrobial activity over PA in solid media, achieving complete ICE25 growth inhibition (10-12 µg/mL) and reduced colony sizes in both strains.

### 3.4. FAs are non-irritant to RHE

FA cytotoxicity was accessed in a keratinocyte cell monolayer to define FAs toxic cell concentration. Irritation effect of defined FAs concentrations was tested on the RHE model. Cell viability of HaCaT cells was reduced in the presence of LA, PA and LIN, except LA at 1/2 MIC for ICE25 (7.81 µg/mL) (Figure 5A). However, 3D RHE models demonstrated no significant viability reduction with ethanol (5%) or any tested FA concentrations (LA-7.81 µg/mL, PA-31.25 µg/mL, LIN-250 µg/mL) (Figure 5B). Morphological analysis revealed slight stratum corneum thickening in LA-treated RHE compared to untreated controls (Figure 6A and B), likely due to LA’s occlusive and hydrating effects, associated with its interaction with stratum corneum lipid bilayers, which loosen their packing and/or fluidize the lipid chains [20]. PBS controls caused stratum corneum desquamation due to its hydrophilic nature favouring occlusion over penetration (Figure 6B). These findings confirm that 7.81 µg/mL LA has no negative impact on RHE, supporting the low cytotoxicity observed in MTT assays (Figure 5B).

**Figure 5.**
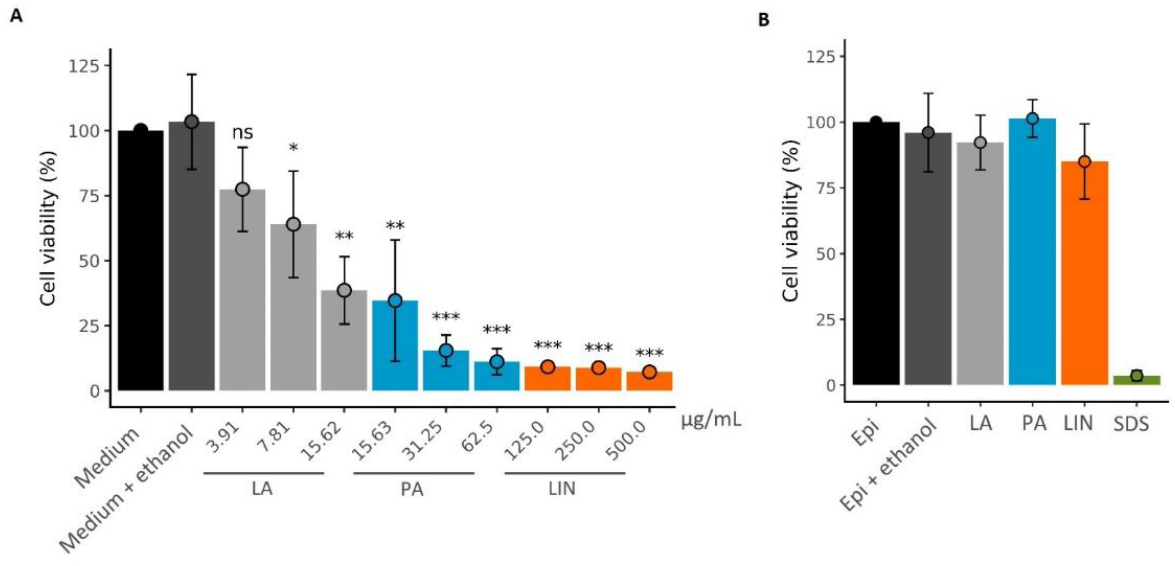
Keratinocytes HaCaT and RHE viability in the presence of LA, PA and LIN. For keratinocytes viability was tested at MIC/2, MIC and 2MIC for each FA, while RHE viability was evaluated at the determined MICs (LA 7.81 µg/mL, PA 31.25 µg/mL and LIN 250 µg/mL). Values are represented as the average ± standard deviation (n=3). t-test results are represented for comparison of positive control (medium) and each one of the conditions. ***p-value < 0.001; **p-value < 0.01, *p-value < 0.05.

**Figure 6.**
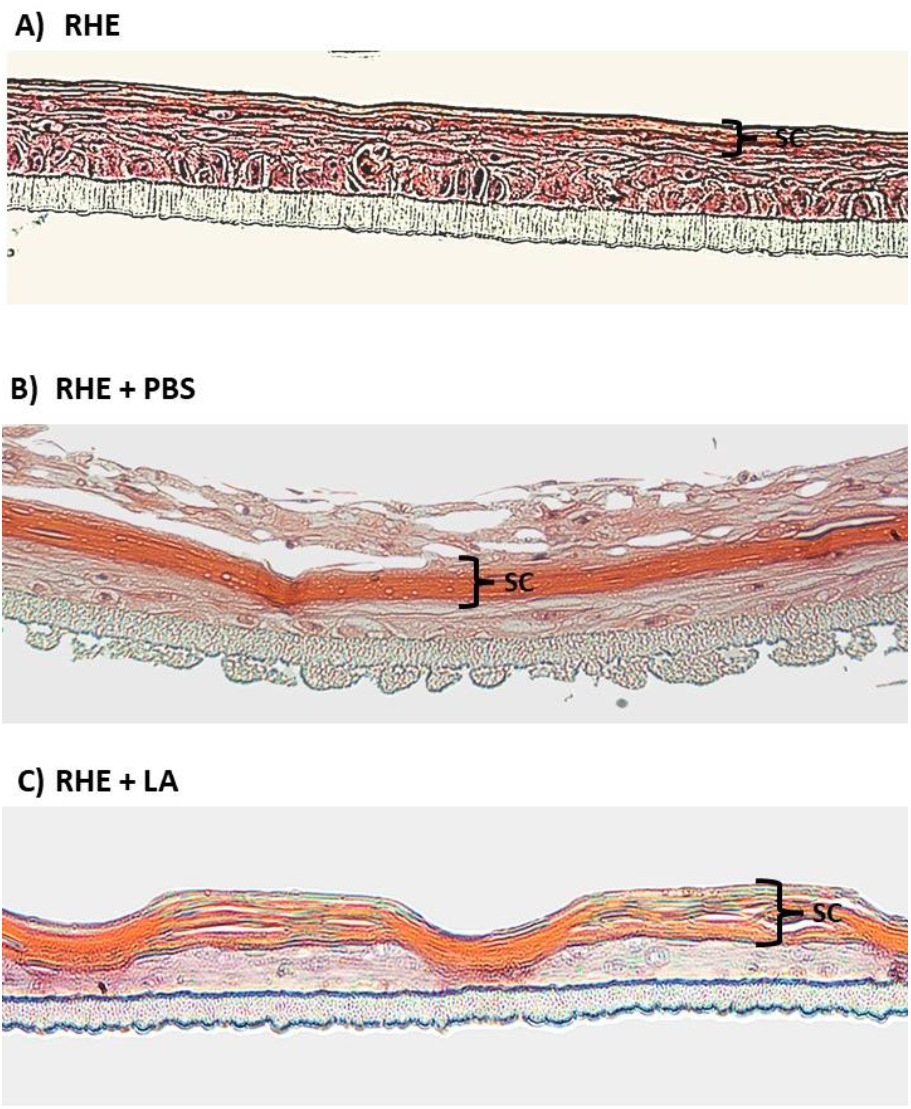
RHE model cross-sections histology. RHE slices were stained with hematoxylin–eosin. RHE were fixed after 1 h incubation with 7.81 µg/mL LA. Images for RHE without treatment (A) and in the presence of PBS (B), used as controls, are also presented. (SC – stratum corneum)

### 3.5. LA tend to reduce ICE25 population in an RHE model

Since LA showed higher antimicrobial activity against ICE25 than towards 19N, we proceeded with the evaluation of the effect of LA on both SE strains after RHE infection/colonisation. Following RHE colonization and 1-hour treatment with 7.81 µg/mL LA, growth reduction was 9±42% for 19N and 29±20% for ICE25, with near-significance for ICE25 versus control (p=0.055) (Figure S2). High standard deviations reflect CFU counting variability in LA presence, consistent with other assays. Although, RHE infected with either strain showed no morphological changes upon LA application (Data not shown).

## 4. Discussion

Endogenous lipids, particularly free FA, exhibit potent antimicrobial activity against both gram-positive and gram-negative bacteria [21]. The FAs selected for this study, LA and PA, have demonstrated antimicrobial effects against several SE strains [11,13,22], whilst LIN has shown differential antimicrobial activity amongst *S. aureus* strains [23]. Here, we evaluated the antimicrobial activity of these FAs against representative strains from lineage B (19N) and lineage A/C (ICE25).

### 4.1 Differential susceptibility between SE lineages

MIC assays in liquid media revealed two key findings: 1) ICE25 showed greater susceptibility to all tested FAs compared to 19N, consistent with hypotheses based on genomic specificities of these lineages [7,16,17], and 2) SE adaptation to FA presence followed the order LA > PA > LIN. The stronger inhibitory effect of LA against SE has been observed previously [11], indicating that, for SE, saturated medium-chain FAs possess higher antimicrobial efficacy against SE.

It was previously described that SE has a lower susceptibility to LIN relative to *S. aureus* planktonic cells [16] and that SE produces less biofilm than *S. aureus* [13]. SE resistance to LIN may be associated with the presence in the SE genome of homologs of the FA inducible efflux pump (FarE) present in LIN resistant *S. aureus* strain [24]. Notably, only 19N possesses a farE homologue associated with endogenous FA efflux, which may explain ICE25’s lower resistance to all three tested FAs. Furthermore, *cis*-9 FAs such as LIN can be detoxified by *S. aureus* oleate hydratase (ohyA) [25]. Our experiments demonstrated that LIN had weak effects on both strains, with ICE25 appearing more susceptible based on MIC assays. This may be attributed to 19N possessing two copies of the OhyA gene compared to ICE25’s single copy, thus providing 19N with greater LIN detoxification potential.

Another resistance mechanism involves induction of the T7SS virulence factor export system upon exposure to LIN and other *cis*-unsaturated FAs in *S. aureus* [26]. In SE, the T7SS genes encoding membrane-associated proteins are found only in the B lineage [7], corroborating 19N’s higher resistance to LIN. Additionally, lineage B strains may exhibit greater FA resistance due to staphyloxanthin biosynthetic genes (crtOPQMN) [27], with LIN stress inducing up-regulation of *crt* genes in *S. aureus* [23]. Staphyloxanthin, a carotenoid anchored to the cell membrane, only present in 19N, can contribute to stabilising membrane fluidity.

The differential SE sensitivity to FAs may also relate to their predominant ionization forms at experimental pH. The ionization constants, pKa, of free LA, PA, and LIN are 5.3, 5.0, and 5.2, respectively [28]. At pH 5.5, all three FAs exist in both protonated and deprotonated forms. SE cells possess a thick peptidoglycan layer interspersed with teichoic acids, conferring an overall negative charge to the cell wall. Thus, all three FAs should interact with cell wall components, with ionization state not explaining differential growth inhibition. However, higher FA concentrations, particularly for long-chain FAs (PA and LIN), could lead to partial monolayer formation around bacterial cells, contributing to increased FA pKa. Kanicky et al (2002) reported that at 0.66 M LIN concentration (approximately sixty-fold higher than our maximum concentration), pKa=9.24, with most of the FA molecules non-protonated at pH 5.5 [28]. This condition promotes stronger FA-bacterial cell wall interaction, potentially explaining effects observed at higher LIN concentrations with possible impacts on cell division.

### 4.2. Growth kinetics and stress adaptation

LA and PA presence in TSB increased lag phase duration, reduced the exponential growth rates and decreased maximum populations, with lag phase effects more pronounced in 19N. Extended lag phases have been reported previously for *S. aureus* in PA and LIN [29]. Prolonged adaptation periods to FA presence suggest their action as cell stressors, with SE cells initiating division only after extended periods, likely due to cell wall and membrane rearrangements [30]. Increased lag time represents a common bacterial stress adaptation trait associated with the metabolic shifts necessary for stress tolerance [31,32]. In *S. aureus* at pH 5, second-generation times are shorter than first-generation times, suggesting active division commences when metabolic processes adapt to LA stress [33]. Following this rationale, the observed lag phase increase in 19N could be explained by FarE biosynthesis induction (see above).

Exponential growth rate and maximum population were significantly reduced with increasing PA concentrations only for 19N, suggesting that ICE25 division requires higher PA concentrations. Conversely, the LA impact was greater for ICE25. This differential behaviour between strains and FAs may be related to aliphatic chain length and saturation or the FA concentration threshold required to induce coping mechanisms.

### 4.3. Solid media growth and colony size

LA demonstrated antimicrobial activity against both strains on agar media, with complete ICE25 growth inhibition at lower concentrations than 19N, consistent with liquid media patterns. For ICE25, a slightly lower MIC was obtained relative to TSB growth. Both strains showed reduced colony radii in LA presence, with 19N particularly affected by PA. Smaller colony sizes may result from slower growth rates, delayed initiation, or early growth cessation [34]. Indeed, lower growth rates and/or extended lag phases were determined for each strain in TSB with LA and PA presence. However, the colony size may be influenced by other factors, such as the production of extracellular polysaccharides, which tends to change colony morphology and increase its radius [35].

LA induced two predominant SE colony radius populations, a phenomenon observed in staphylococci under stress conditions [34]. This trait may be associated with bacterial subpopulations exhibiting extended lag times, behaving as persisters with increased antibiotic resistance [34], consistent with 19N’s greater colony size reduction compared to ICE25. This differential effect could also be related to the production of extracellular polysaccharides by ICE25. PA induced less heterogeneous colony size distributions than LA, suggesting different antimicrobial mechanisms and distinct SE adaptation strategies for these FAs.

### 4.5. Skin cytotoxicity and compatibility

Keratinocyte cell line assays demonstrated a strong cytotoxic effect, particularly for LIN and PA, whilst no FAs tested showed cytotoxic effect in the RHE model. Sebocytes, multifaceted epithelial cells, viability was unaffected by LA (3.9 μg/mL) [36], highlighting these FAs potential for topical application. In recent years, RHE has been used to assess cytotoxic effects of various molecules and skin treatments [37,38]. Our results underscore the importance of utilising RHE models, as their 3D structure and cell organisation mimicking human skin morphology confer resistance to FA cytotoxicity. Since the generated RHE model does not include sebaceous glands, the study of a possible synergic antibacterial effect of tested FAs with endogenous FAs was not discussed.

Testing FAs MIC concentration, resulted in tissue viabilities above 50% (Figure 5) and were classified as non-irritants. Additionally, increased stratum corneum thickness was observed in non-infected RHE treated with LA compared to PBS controls. The epidermis is the uppermost layer of the skin, generated by keratinocyte differentiation, consisting of the viable epidermis and the non-viable external layer, stratum corneum, which is in contact with the environment. The SC is commonly presented schematically as “brick and mortar” model, in which differentiated non-viable keratinocytes (corneocytes) are embedded in a lipid matrix [39]. Free FAs are important components for SC hydration maintenance and intercellular lipid barrier establishment. LA skin layer penetration has been confirmed previously [40] and corroborated by our histological results.

### 4.6. RHE infection model and therapeutic potential

Recent studies have employed RHE models to test the antimicrobial effects of various compounds against *Staphylococcus* species [38]. No visible morphological changes were observed in RHE infected with either strain following LA application, possibly attributable to relatively short exposure time (1 hour) post-bacterial infection. This highlights the need for further investigation regarding LA impact over extended exposure times, various concentrations and different infection time points. The application of LA at the beginning of RHE infection, would allow to test its potential protective effect.

Our RHE infection model results align with MIC and growth parameter determinations: ICE25 demonstrated greater LA sensitivity compared to 19N, with three-fold higher CFU reduction following LA treatment. These results suggest LA topical application could prevent pathogenic SE proliferation at the skin level, thus minimizing potential infection development whilst preserving beneficial commensal strains.

## 5. Conclusions

This study aimed to assess whether selected FAs exhibit discriminative antimicrobial activity against *SE* strains belonging to the A/C and B lineages, intending to develop a topical product for selective disinfection. We demonstrated that FA (LA and PA) induced a decrease in maximum population and reduced growth rates, with effects more pronounced for the ICE25 strain (A/C lineage), whilst extended lag phases predominantly affected 19N (B lineage). In solid media, greater colony size reduction in LA presence suggests negative growth impacts, that through persister populations may contribute to reduced antimicrobial efficacy. Nevertheless, ICE25 showed a three-fold higher percentage of CFU reduction compared to the B lineage strain.

Collectively, our results demonstrate that endogenous FA (LA, PA and LIN) exhibited greater antimicrobial activity against A/C lineage than B lineage strain representatives. Literature evidence and genome comparisons between strains suggest that molecular responses of A/C and B SE strains to FA presence differ and are FA-dependent. These findings, combined with the determined low FA cytotoxicity, support the use of LA as a selective topical antimicrobial specifically targeting SE strains associated with infection whilst preserving beneficial commensal populations.

## Declarations

### Ethics approval and consent to participate

Not applicable

### Consent for publication

Not applicable

### Availability of data and materials

All data generated or analysed during this study are included in this published article, its supplementary information files and through https://github.com/eccmorais/MIC_Fatty_acids_Sepidermidis.git

### Authors’ contributions

E.M., D.P.C.B., L.G.G., A.O., M.M. and A.V.C. contributed to the study conception and design. E.M., F.M. and O.B. performed material preparation, data collection and analysis. All authors were involved in data interpretation. E.M. wrote the first draft of the manuscript, and all authors commented on and contributed to the manuscript’s writing. All authors reviewed the manuscript.

## Acknowledgements

Not applicable

