## Supplementary Materials and Methods for "Lauric acid: a promising antimicrobial for the selective inhibition of *Staphylococcus epidermidis* strains associated with infection"

**Study design**

The antimicrobial effects of three FAs (LA, LIN and PA) were tested in different growth conditions: in liquid media, in agar and on a 3D reconstructed human epidermis (3D-RHE) model. MIC values were determined in liquid and solid media, TSB was also used to trace the growth curves and RHE to better mimic *in vivo* like skin conditions in FA skin irritation and inhibition of SE strains tests. As a key component of the epidermis, keratinocytes are crucial in the initial protective skin response; they play an active part in the immune response, inflammatory processes, and wound healing [1]. Cell viability was used to test the cytotoxic effect of FAs. The *in vitro* skin irritation test on RHE was performed according to the conditions established in OECD Test Guideline 439 [2]. RHE, consisting of human keratinocytes, mimics the native human epidermis in terms of morphology and lipid composition [3]. The antibacterial effect of nontoxic FAs concentrations on bacterial strains was further tested on RHE by histology studies.

**Bacterial strains and inoculum preparation**

For this study, we selected representative strains of SE phylogenetic lineages B and A/C, respectively, 19N collected from the anterior nares of a healthy person and ICE25 isolated from the respiratory secretions of a patient [4]. All bacterial cultures were grown at 37°C in tryptic soy agar plate (TSA, BactoTM, pH 7.4). A single colony was then picked and streaked in a TSA plate at pH 5.5, adjusted with HCl 37% before sterilization, and incubated overnight. For the assays in tryptic soy broth medium (TSB, BactoTM), minimum inhibitory concentration (MIC) and growth curve determinations, a single colony was inoculated overnight in TSB at pH 5.5 with 180 rpm. For the experiments in TSA, one colony was picked from the TSA plate at pH 5.5 and streaked again in a new plate. All bacterial assays in this study were carried out at pH 5.5.

**Preparation of FA stock solutions**

FAs selected based on reported antimicrobial activity against SE were used in this study: lauric acid (LA, C12:0, TCI Chemicals), palmitoleic acid (PA, C16:1, Acros Organics) and linoleic acid (LIN, C18:2, ALFA AESAR) (Table S1). A stock solution was prepared in absolute ethanol and then diluted in TSB or TSA media supplemented with 0.1% (v/v) of DMSO, a concentration which is not toxic to SE [5]. For cytotoxicity assays and assays with RHE, FA stock solutions were diluted in phosphate-buffered saline (PBS). The final ethanol concentration was kept at 0.5% (v/v) in all assays, except for cytotoxicity assays for which the ethanol concentration was 5%. FA stock solutions were always prepared on the assay day.

**Determination of Minimum Inhibitory Concentration (MIC) of FAs**

The broth microdilution method with TSB was used to determine the fatty acids' MICs for three biological replicates (independent pre-inoculum) per experimental condition. To prepare the inoculum, the overnight pre-inoculum in TSB was adjusted to OD_600nm_ of 0.06 (aprox. 1.5 x 10^8^ CFU/mL), and further dilutions were carried out to obtain an initial inoculum of 5x10^5^ cells in each microplate well (final volume 200 µL). As previously described, the TSB medium was supplemented with fatty acids to obtain the desired concentrations (Table S1). After 20 hours of incubation at 37°C, optical density at 595 nm (OD_595nm_) was measured in a microplate reader (Infinite ®200 PRO series, Tecan Group Ltd., Mannedorf, Switzerland) and the percentage of growth was calculated considering the OD_595nm_ in the control condition (${OD}_{0}$, in the absence of FA and in the presence of 0,1% (v/v) DMSO and 0,5% (v/v) ethanol) as 100%:

$Growth (\%) =\frac{{OD}_{FA}}{{OD}_{0}}*100$	(1)

where ${OD}_{FA}$ corresponds to OD_595nm_ when TSB was supplemented with FAs.

**Growth curve determination in TSB supplemented with FAs**

Growth curve determinations were performed for FA sub-inhibitory concentrations for three biological replicates (Table S1). In the control condition, without FA, 0,1% (v/v) DMSO and 0,5% (v/v) ethanol were added. To prepare the inoculum, the overnight pre-inoculum in TSB was used to obtain an initial OD600nm of 0.06. This growth assay was performed with 100 mL of TSB (in 250 mL Erlenmeyer) in aeration conditions (225 rpm, Brunswick Innova 4300 Incubator Shaker). Bacterial growth was followed by measuring OD_600nm_ every hour, until the stationary phase.

Each growth curve was fitted with the ‘logisticlag1’ model from the Curveball Python Library [6]. This model includes a lag phase to better fit growth data. The ‘logisticlag1’ model defines the population size (N) by the following reaction:

$N(t)= \frac{K}{\left[ 1-\left( 1-\left( \frac{K}{N_{0}} \right)^{v} \right)e^{-rvA(t)} \right]^{1/v}}$, where $A(t)$ is defined as: $A(t)= t + \frac{1}{V}log\left( \frac{e^{-Vt} + q0}{1 + q0} \right)$	(2)

Where $N_{0}$ is the initial population size, $K$ the maximum population, $r$ the initial per capita growth rate; $v$ the curvature of the logistic term, $t$ the time, $V$ the adjustment rate and $q0$ the initial adjustment to the current environment. After model fitting, lag time and exponential growth rate were estimated using curveball.models.find_lag and curveball.models.find_max_growth functions.

**SE growth in TSA supplemented with FAs**

To assess the effect of FAs on SE growth in solid media, 0.5 x 10^3^ CFUs were plated onto TSA at pH 5.5 with several FA concentrations (Table S1). After 24 hours of incubation, each plate was observed under the on-axis zoom microscope Axio Zoom.V16, and an image was captured from each of five random plate regions with an incorporated Zeiss Axiocam 503 mono CCD camera controlled with the Zeiss Zen Blue 2.1 software using the 1x8 NA objective under the Bright Field optics. The radius of five colonies was measured for each image considering the colonies' shape as a circle using the software Zeiss Zen Lite Blue 3.8. Bacterial viability was assessed by counting the number of CFUs after 40 hours of incubation, since after 24 hours only a small fraction of the colonies was detectable. The percentage of growth was determined considering the number of colonies in the control condition (${CFU}_{0}$, in the absence of FA and in the presence of 0,1% (v/v) DMSO and 0,5% (v/v) ethanol) as 100% (Equation 3):

$Growth (\%) =\frac{{CFU}_{FA}}{{CFU}_{0}}*100$	(3)

where ${CFU}_{FA}$ corresponds to the number of CFU counted for each plate supplemented with FAs.

**Cytotoxicity assay on 2D monolayer**

Immortalized human keratinocytes, HaCaT, were cultured in Dulbecco’s Modified Eagle’s Medium (DMEM) supplemented with 10% fetal bovine serum (FBS) and 0.1% Pen Strep (10.000 U/mL penicillin, 10 ug/mL streptomycin). Cells were maintained at 37°C in a 5% CO_2_ humidified atmosphere and after 24h media was exchanged to remove the residual solvent (dimethyl sulfoxide, DMSO) from the cryo-conservation medium. Cytotoxicity assays were performed by harvesting cells from the flasks at 80% confluence. Cytotoxicity of different concentrations of FAs to HaCaT cells was accessed using the 3-(4,5-dimethylthiazol-2-yl)-2,5-diphenyltetrazolium bromide (MTT) assay. Cells were detached using Trypsin-EDTA (0.25%), and afterward re-suspended in HaCat-supplemented medium to stop trypsin activity. Cell suspensions were centrifuged at 2840 rpm, for 5 min, 20°C. Then were seeded in each well of 96-well plates at a density of 2 x 10^4^ cells/well (per 200 µL) and grown for 24h at 37°C in 5% CO_2_ humidified atmosphere until being in a subconfluence state. Media was replaced by 200µl of DMEM, DMEM with 5% of ethanol, and selected FAs concentrations (MIC/2, MIC and 2MIC) in DMEM/5% ethanol (Table S1). After 3 h of exposure, 10 µL of the 12 mM MTT was added to each well and incubated in a 37°C in 5% CO_2_ humidified atmosphere for 1 h and 45 min. After this, the MTT solution was removed and the cells were washed with PBS. The formazan crystals were dissolved with DMSO for 10 min. The absorbance was measured at 540 nm in a multi-well plate reader (SpectraMax 340PC Microplate Reader, Molecular Devices, LLC., San Jose, CA, USA) and the cell viability was calculated (Equation 4). The negative control (NC) consisted of a cell growth medium (DMEM), defining 100% of cell viability. The assays were performed for 3 biological replicates.

$Cell viability (\%) =\frac{Absorvance of treated cells}{Absorvance of negative control}*100$	(4)

**Reconstructed human epidermis (RHE) generation**

The development of RHE on cell culture inserts with polycarbonate membranes (Merck, Millipore) was performed following the established procedure [3]. For the development of RHE, neonatal human epidermal keratinocytes (HEKn) (passage 3) were cultured in EpiLife^TM^ (Termofisher) medium supplemented with 1% Pen-Strep, 1% Human Keratinocyte Growth Supplement (HKGS), and 0.06 mM CaCl_2_. The medium was changed after one day to remove residual DMSO. Cells were maintained at 37°C in 5% CO_2_ humidified atmosphere until harvested at 70% confluence. After trypsinization, EpiLife medium (8 mL) with 1% HGKS, 1% PenStrep, 0.06 mM CaCl_2_, and 2% FBS were added to block the action of trypsin. Cells were then washed and centrifuged at 2840 rpm, 5 min, 20°C. Afterwards cells were seeded in polycarbonate filter (insert) of 6-well plates at a density of 3.5x10^5^ cells/mL in 500 µL of cell suspension. To each well, 2.5 mL of EpiLife medium with 1% HGKS, 1% PenStrep, and 1.5 mM calcium were added. After 1.5h of incubation, the medium was aspirated from the insert, and at the air-liquid interface the medium of the well was replaced with 1.5 mL of EpiLife medium with 1% HGKS, 1% PenStrep, 1.5 mM calcium, 1% keratinocytes growth factor (KGF), and 100 µg/mL vitamin C. Media of the wells were changed every two days until 12 days when the RHE was morphologically fully differentiated. RHEs were always maintained at 37°C in a 5% CO_2_ humidified atmosphere before further use.

**Cytotoxicity assay on 3D RHE**

To evaluate the cytotoxicity on the RHE model the assay was based on the irritation test described previously [7], and according to the conditions established in OECD Test Guideline 439 [8]. Briefly, 200 µL of the FAs solution (Table S1) were placed above the epidermis formed inside the insert. The following controls were used: 5% SDS as positive control and EpiLife medium with 5% ethanol as negative control. After 3 h of incubation, the FA solution was aspirated, and inserts were washed 3 times with PBS. Then, inside the insert was placed 1 mL of EpiLife medium, with antibiotic and 10% of MTT solution (5 µg/mL). After 3 h of incubation, the MTT solution was removed, and the inserts were cleaned 3x with PBS. Inserts were placed under 2 mL of isopropanol and were left overnight at room temperature. Then, the inserts were punctured to let the solution run into the wells. The cells were resuspended and 200 µL were placed in triplicate into the 96-well plate. The absorbance was measured at 570 nm in the plate reader (SpectraMax 340PC Microplate Reader, Molecular Devices, LLC., San Jose, CA, USA) and cell viability was calculated according to Equation 4.

**Antibacterial studies using RHE**

Upon RHE formation, it was infected/colonized by SE strains. Briefly, 1 x 10^2^ CFUs of 19N and ICE25 strains resuspended in PBS were inoculated on top of each RHE. After 3 h of incubation, the excess of inoculum was carefully removed through decantation, in order to evaluate the effect of LA on the skin adherent bacteria. A total of 100 µL of LA (7.81 µg/mL with 0.5% ethanol) was placed topically on the RHE. For the negative control, 100 µL of PBS with 0.5% ethanol was used. One hour later, 100 µL of the suspension was recovered after up and down pipetting. RHE was washed twice with 100 µL PBS and the recovered volumes were added to the previously recovered suspension. A volume of 100 µL of this suspension was plated onto TSA plates (pH 7.4) to count CFUs. Growth (%) was determined using Equation 3 and considering the CFUs counted on the negative control as corresponding to 100% of growth.

**Histological analyses**

Tissue staining and histological analyses were conducted according to the established method [9]. For the histological analysis, inserts with generated RHE were fixed immediately in 10% neutral-buffered formalin (Sigma- Aldrich, St. Louis, MO, USA) for a minimum of 24 h at room temperature. Then, the membrane of the insert containing the RHE was removed and kept in formalin until the fixation procedure. Skin sections (5 mm thick) were mounted on slides for histological analysis. Tissue sections were stained with hematoxylin–eosin to allow a standard morphological analysis of the RHE. Images were obtained using the Nikon Eclipse TE2000-S fluorescence microscope (Nikon instruments, Melville, NY, USA) and analyzed with the ImageJ Software version 1.53q.

**Statistical analysis**

Two comparisons were made for the several assays: 1) between strains for a certain experimental condition; and 2) within the 19N/ICE25 strains against the respective controls (0 µg/mL FAs). *T*-tests were used for MIC, growth in TSB, and TSA-related data. For colony size analyses in the presence of FAs Mood’s median test was performed using SciPy (Virtanen, 2020). Results were considered statistically significant if a p-value <0.05 was obtained in any of those statistical tests.

All data analyses described were performed in Python 3.11.5 using in-house scripts and are available through GitHub access at <https://github.com/eccmorais/MIC_Fatty_acids_Sepidermidis.git>.

[8]	Test Guideline No. 439: In Vitro Skin Irritation: Reconstructed Human Epidermis Test Methods. 2021.

[9]	de Barros DPC, Santos R, Reed P, Fonseca LP, Oliva A. Design of Quercetin-Loaded Natural Oil-Based Nanostructured Lipid Carriers for the Treatment of Bacterial Skin Infections. Molecules 2022;27. https://doi.org/10.3390/molecules27248818.
